# Identification of transcriptional signatures for cell types from single-cell RNA-Seq

**DOI:** 10.1101/258566

**Authors:** Vasilis Ntranos, Lynn Yi, Páll Melsted, Lior Pachter

## Abstract

Single-cell RNA-Seq makes it possible to characterize the transcriptomes of cell types and identify their transcriptional signatures via differential analysis. We present a fast and accurate method for discriminating cell types that takes advantage of the large numbers of cells that are assayed. When applied to transcript compatibility counts obtained via pseudoalignment, our approach provides a quantification-free analysis of 3’ single-cell RNA-Seq that can identify previously undetectable marker genes.

---

Single-cell RNA-Seq (scRNA-Seq) technology provides transcriptomic measurements at single-cell resolution, making possible the identification and characterization of cell types in heterogeneous tissue. The problem of identifying genes that are differential between cell groups is analogous to the differential expression problem in bulk RNA-Seq. Bulk RNA-Seq differential expression methods can be applied directly to test genes for differences between groups of cells^1^, and methods that account for technical artifacts in scRNA-Seq experiments such as by modeling dropout seem to offer some advantages.^2,3^ However, one aspect of scRNA-Seq that current methods do not take advantage of is the large number of cells sampled in single-cell experiments.

In this paper, we show how prediction methods that take advantage of large numbers of cells can greatly improve gene-level differential expression results. We focus on logistic regression, which was considered when microarray gene expression assays were developed^4,5^, but abandoned due to limited sample sizes. scRNA-Seq provides the large number of samples (i.e. single cells) required to accurately fit a logistic regression model. Conversely, fitting a linear logistic regression is near linear in the number of cells, which is imperative in the era of growing scRNA-Seq experiments. Instead of the traditional approach of using the cell labels as covariates for gene expression, we perform logistic regression for each gene to predict cell labels from the quantifications of constituent transcripts. Fitting this model provides the optimal linear combination of transcript quantifications that distinguishes cell groups, providing information about effect sizes of constituent transcripts, i.e. the “direction of change” (Figure 1a—h). Unlike traditional methods that test either for changes in overall gene abundance or for differential transcript usage, our method provides a unified testing framework that eliminates the need for such a dichotomy (Supplementary Figure 1).

**Figure 1.**
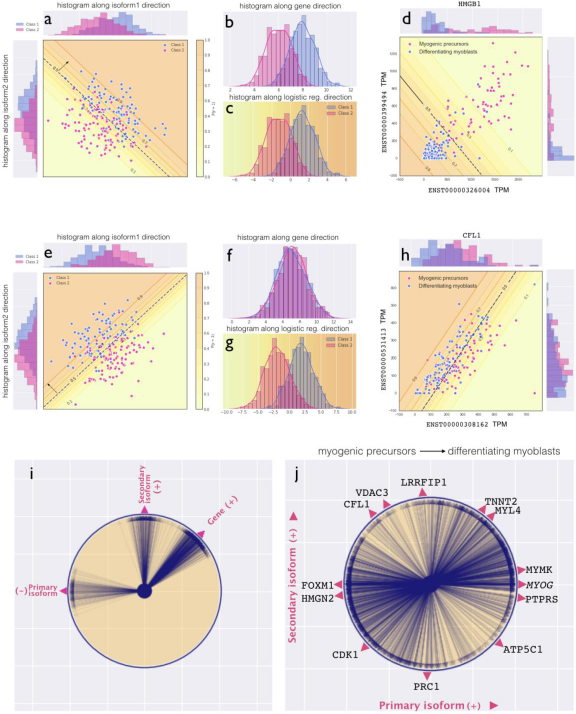
Logistic regression applied to scRNA-Seq. Logistic regression can be used to detect gene differential expression at isoform level resolution. Panel **(a)** shows a hypothetical scenario with two cell groups (s‘Classl’ and ‘Class 2’, colored blue and pink respectively) where both isoforms change with the same effect size reflecting gene overexpression. In this case the direction of change discovered by logistic regression (dashed line) projects the points along the [1,1] vector. This operation **(c)** is equivalent to the conventional approach of summing the corresponding isoform counts and directly testing for gene differential expression (‘the gene direction’) **(b)**. Panels **(b)** and **(c)** show histograms comparing abundances projected to the gene direction and the direction learned by logistic regression respectively. Panel **(d)** shows gene HMGB1 from the Trapnell *et al.* data set that demonstrates similar behavior during myoblast differentiation. Panel **(e)** considers the case where the two isoforms have opposite effect sizes (isoform switching) that cancel each other out when projected along the gene direction. Even though gene counts cannot distinguish between the two populations **(f)**, logistic regression is able to learn that a difference in isoform abundances is best for distinguishing the classes, thus effectively detecting the isoform switching event **(g)**. Panel **(h)** shows gene CFL1 from the Trapnell *et al.* data set demonstrating similar behavior during myoblast differentiation. Panel **(i)** shows a simulation where the “directions of change” in 1000 genes with two isoforms were randomly chosen to be either in the gene direction or along each isoform individually (see Methods). The circle plot overlays the directions detected by logistic regression for each gene. Panel **(j)** shows the directions of change of 1308 genes from the Trapnell *et al.* data set that were identified by logistic regression as differentially expressed between myogenic precursors and differentiating myoblasts (see Methods). The direction of the arrow indicates the change in expression of a gene *from* myogenic precursors *to* differentiating myoblasts. For example, in CDK1, where the expression of both isoforms decreases from myogenic precursors to differentiating myoblasts, the corresponding arrow points to the southwest.

In a simulation based on experimental effect sizes (see Methods), logistic regression outperforms other existing scRNA-Seq gene differential expression methods, even with different normalizations (Supplementary Figures 2, 3b). The advantage comes from the ability of logistic regression to identify the optimal linear combination of isoforms for differential analysis. In the case that isoforms move in concert, naïve gene quantification by summing of isoform counts performs similarly to logistic regression (Figure 1b,c and Supplementary Figure 2c,d), but in the event of isoform switching logistic regression has a substantial advantage (Figure 1f,g and Supplementary Figure 2e,f). When applied to a data set of differentiating myoblasts from Trapnell *et al*.,^6^ the method reveals the nature of transcript dynamics across multiple genes known to be important for myogenesis (Figure 1i,j). We found a diversity of transcript dynamics, suggesting that methods that test only for changes in overall gene expression or only for differential transcript usage are likely to miss a significant proportion of our differential genes (Supplementary Figure 1b).

**Figure 2.**
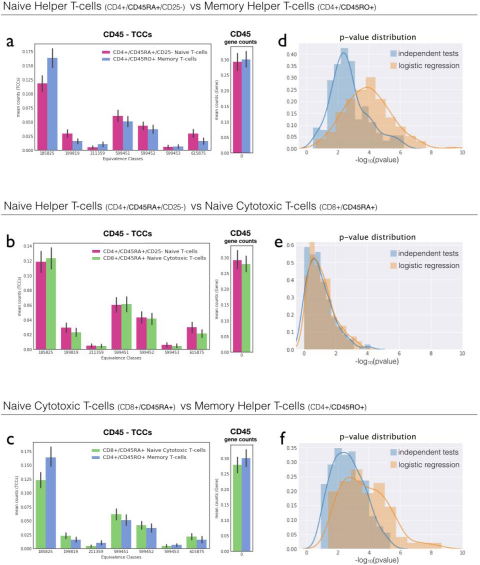
Logistic regression discovers CD45 in purified T-cell types. Logistic regression was used to perform pairwise differential expression analysis of purified memory T-cells, naïve T-cells, and naïve cytotoxic T-cells that were sequenced with 10X. In **(a, c)**, logistic regression on TCCs recovered the separation between CD45RA+ and CD45RO+ T-cells, while summing counts to produce gene abundances masks the differential expression. Furthermore, independently testing the TCCs and then performing Bonferroni correction reduces power **(d, f)**. In contrast, when testing CD45RA+ naïve helper T-cells and CD45RA+ naïve cytotoxic T-cells **(b, e)**, CD45 was not significant using logistic regression or gene counts **(b)**, and there was little difference in p-value distribution between independently testing the TCCs and performing logistic regression **(e)**.

While transcript quantifications are biologically meaningful, in some cases they may be infeasible to obtain.^7, 8^ We therefore examined the possibility of performing logistic regression directly on the transcript compatibility counts (TCCs) obtained via pseudoalignment.^9^ TCCs, which were introduced in Ntranos *et al.^10^* as model-free transcriptomic signatures for single cell clustering, constitute the sufficient statistics needed for quantification. By virtue of being unprocessed, they represent the “raw data” better than quantified gene expression profiles.^11^ On simulated data, the performance of logistic regression with TCCs is similar to that with transcript quantifications (Supplementary Figure 3a). Another advantage of TCCs is that they are readily computable from 3’ capture and sequencing single-cell data. To investigate whether logistic regression with TCCs confers an advantage over gene-count based differential analysis from such data, we examined 10X Chromium scRNA-seq from three T-cell populations that were purified using antibodies specific to different isoforms of CD45 (PTPRC).^7^

Using TCCs, we performed pairwise differential analyses of purified CD45RO+ memory helper T-cells, CD45RA+ naïve helper T-cells, and CD45RA+ naïve cytotoxic T-cells, providing two positive controls (CD45RA+ vs CD45RO+) and a negative control (CD45RA+ vs CD45RA+) for the method. Logistic regression was able to detect differential expression of CD45 in the purified CD45RO+ memory and CD45RA+ naïve T-cell populations (Figure 2a,c). This result was deemed impossible in Peterson *et al*.,^12^ where it was noted that 3’ mRNA sequencing alone could not resolve these markers. We confirmed that gene counts alone cannot identify CD45 as differential (Figure 2a-c), and furthermore found that independent testing of TCCs reduced statistical power (Figure 2d—f). The two CD45RA+ naïve T-cell populations had similar TCCs as expected, and CD45 was not identified as differential between them (Figure 2b). Further examination of the transcripts corresponding to the equivalence classes identified by logistic regression pointed at which transcripts were differentially regulated (Supplementary Figure 4). Visual inspection of the differential equivalence classes identified by logistic regression for CD45 revealed that the corresponding isoforms were being distinguished by virtue of alternative unannotated 3’ untranslated regions (UTRs) (Supplementary Figure 4). To quantify the extent of isoform accessibility by 3’-end sequencing, we estimated the distribution of read pseudoalignments with respect to the annotated 3’UTRs (Supplementary Figure 5). We found a substantial number of reads distal to annotated 3’-ends, pointing to a large number of unannotated 3’ UTRs. The results on CD45 are concordant with previous work on lymphocytic surface receptor isoform diversity^13^. Equivalence classes provide access to isoforms in genes other than CD45; we found multiple other genes that also exhibited isoform switching between memory and naïve T cells (Supplementary Figure 6).

To examine whether TCCs are informative in a *de novo* scRNA-Seq experiment, we analyzed the PBMC dataset,^7^ which consists of 68579 cells sequenced at an average of 20491 reads per cell. After clustering and using known cell type markers to annotate the clusters (Supplementary Figure 7), we were able to recapitulate our previous CD45 differential analysis: CD45 was identified as differential between memory and naïve T-cells respectively (Supplementary Figure 8), showing that TCC-based logistic regression can be applied to cell groups generated by unsupervised clustering as in standard, *de novo* scRNA-Seq workflows.

While logistic regression is a simple method, it is especially powerful for scRNA-Seq since it leverages the large number of cells available in scRNA-Seq experiments and incorporates isoform-information for gene-level testing. Logistic regression reveals the contribution of individual isoforms to the gene-level differential analysis, aiding in interpretability of results. While we have demonstrated the power of logistic regression for performing gene-level differential expression between two cell types, any two cell groupings can be used. Furthermore, logistic regression can be performed on all genes instead of constituent isoforms of a single gene to discover gene markers characterizing cell types. Finally, our method scales effectively with both the number of reads and cells, which is critical for processing increasingly large scRNA-Seq datasets.

## Methods

### Trapnell *et al.* 2014 analysis

We downloaded the preprocessed Trapnell *et al.* 2014 data from the conquer database,^1^ which included the quantified transcript-per-million (TPM) values and cell labels for 222 serum-induced primary myoblasts over a time course of 0, 24 and 48 hours. We selected the 85 myogenic precursors and the 97 differentiating myoblasts for differential expression analysis. We used Ensembl Homo_sapiens.GRCh38.rel84.cdna.all.fa to group 176241 transcripts into 38694 genes and tested each gene for differential expression between myogenic precursors and differentiating myoblasts using logistic regression over the constituent isoforms. After Benjamini-Hochberg correction, we obtained 1308 significant differential genes (< 0.01 FDR). We visualized these genes in a circular plot by performing logistic regression on the primary and secondary isoforms, which are defined as the isoforms with the largest and second largest average expression over all cells, and plotted the direction of change identified by logistic regression.

### Zheng *et al.* 2017 analysis

We obtained the raw reads for the three human PBMC purified cell sub-type datasets, CD4+/CD45RA+/CD25-naïve T-cells, CD4+/CD45RO+ memory T-cells and CD8+/CD45RA+ Naïve cytotoxic T-cells, from Zheng *et al.,* 2017. The reads were preprocessed (barcode detection, error-correction and pseudoalignment) with the scRNA-Seq-TCC-prep kallisto wrapper (SC3Pv1 chemistry) to obtain the single cell transcript compatibility counts (TCC) matrix (https://github.com/pachterlab/scRNA-Seq-TCC-prep). After filtering out cells with total UMI counts outside the interval [1K-30K], we obtained 31831 cells (9923 CD4+/CD45RA+/CD25-naïve T-cells, 9994 CD4+/CD45RO+ memory T-cells and 11914 CD8+/CD45RA+ naïve cytotoxic T-cells respectively). We selected all the equivalence classes that contained at least one isoform associated with the CD45 gene (a.k.a. PTPRC, ENSG00000081237, ENSG00000262418) and filtered out the ones with total UMI counts less than 0.25% of the total number of cells, i.e. equivalence classes with fewer than ~79 UMI counts across all cells. This resulted in 7 equivalence classes uniquely associated with subsets of the annotated isoforms of the CD45 gene. The gene counts for each cell were obtained by summing the corresponding TCCs. We performed all three pairwise tests for differential expression between the purified cell sub-types using a logistic regression model on the 7 TCCs, on the aggregated gene counts, and independently on each equivalence class. For each pairwise test, we randomly subsampled 3000 cells per group across 200 Monte-Carlo iterations to generate error bars and p-value distributions.

The raw reads for the 68k PBMC dataset were preprocessed with the scRNA-Seq-TCC-prep kallisto wrapper to obtain the TCC matrix. Equivalence classes that mapped to multiple Ensembl gene names and cells with total UMI counts outside the interval [2K-20K] were filtered out. The resulting 65444 cell x 95426 equivalence class matrix was subsequently used for post-processing and clustering with scanpy.^14^ We used the same steps outlined in “Zheng et al. recipe” that was included in scanpy, except we selected the 5000 most variable equivalence classes in lieu of selecting the 1000 most variable genes. To verify the clustering structure, we plotted the cells on t-SNE space according to their expression of specific marker genes (obtained by summing all the corresponding TCCs). Supplementary Figure 8 focuses on the clusters that most likely correspond to populations of naïve cytotoxic T-cells (Cluster A, CD8A+/CD4-/CCR7+, n=5226 cells), naïve T-cells (Cluster B, CD4+/CCR7+, n=12424 cells) and memory cytotoxic T-cells (Cluster C, S100A4+/CCR10+, n=4173). Clusters A and C corresponded to clusters 3 and 6 in Supplementary Figure 7, whereas cluster B was obtained by manually merging clusters 1 and 2. Following the same steps as in the purified 10x datasets, we performed all three pairwise tests for DE in the CD45 gene. The bar plots with standard errors were generated by randomly subsampling 2000 cells per cluster across 200 Monte-Carlo iterations. The average p-values were reported.

### Estimation of the read location distribution

In order to estimate the distribution of distances to the 3’ end we pseudoaligned the reads from the three purified T-cell populations to the transcriptome using the pseudobam option of kallisto 0.44. In case of multiple alignments, the weight of one read was split evenly across all reported transcripts. The distance to the 3’ end was inferred from the transcriptome coordinates reported in the BAM file.

### Simulation for Figure 1i

Two isoforms in each group (n=200) for each gene were sampled from *N(μ_1_,I)* and *N(μ_2_,I)* with *μ_1_* =[5,5], *μ_2_* = [5+cos(θ), 5+sin(θ)] respectively and θ was chosen for each gene to be π/4 (increase in gene direction), π/2 (increase in primary isoform direction) or π (decrease in secondary isoform direction), with corresponding probabilities [.6, .25, .15]. The circle plot in Figure 1i overlays the directions of change that were detected from individually fitting a logistic regression model for each gene.

### Simulation framework

scRNA-Seq simulated data was generated by learning features from the scRNA-Seq data in Trapnell *et al.* 2014. In each simulation, cells were simulated from two different cell types: a null type and a perturbed type, each with 105 cells. The null type was modeled after the cluster of proliferating myoblasts from the Trapnell *et al.* 2014 dataset. Specifically, after quantification of the dataset using kallisto and clustering on TCCs, the cluster containing cells with MYOG expression was identified and the resulting TPMs from this cluster were used to estimate the parameters of a lognormal distribution for each transcript. To simulate the null cell type, TPMs for each transcript were drawn from a truncated lognormal distribution. This approach to modeling cell-by-cell variability was motivated by the Tobit model on TPMs used in Monocle.^2^

Three different types of simulated data were prepared to reflect distinct perturbation scenarios and effect sizes. Transcripts expressed in fewer than 5 of the 105 cells were deemed too lowly expressed and filtered out from the perturbation. In the independent effects simulation, 30% of the transcripts that passed the filter (20456 out of 68179 expressed transcripts) were chosen at random to be perturbed. For each transcript, a minimum effect size of 1.5-fold was drawn from a truncated lognormal distribution. The direction of each perturbation was chosen uniformly at random (50% upregulated, 50% downregulated). In the correlated effect simulations, genes with all transcripts passing the filter also passed the filter. 30% of remaining genes (~5220 of 17390 genes) were chosen at random to be perturbed and expressed transcripts (defined as expressed in >= 5 cells) of that gene were perturbed with the same effect size drawn from a truncated log normal distribution at a minimum of 1.5. In the experiment-based simulations, the effect sizes were learned from Trapnell *et al.,* 2014 from the set of transcripts that either DESeq2^15^ or sleuth^16^ found to be differentially expressed. Random genes were chosen to be perturbed for the simulation. Each perturbed gene in the simulation was matched to an experimentally differential gene, and their effect sizes were matched according to the relative abundance of each transcript.

The effect sizes were applied to the mean expression, and abundances per cell were generated by sampling from lognormal distribution truncated at zero. Given these cell-by-cell abundances, RSEM^17^ was used to generated uniformly sequenced paired-reads, using an RSEM model learned from a proliferating myoblast cell from the Trapnell *et al.,* 2014 data set and a background noise read percentage (parameter theta) of 20%. The number of reads per cell is learned from the myoblast cluster by fitting a lognormal distribution of reads per cell (u = 14.42, ro = 0.336), corresponding to a mean of 193,000 paired-end reads per cell, 210 cells per simulation, and 40,530,000 paired-end reads per simulation.

### Simulation analyses

Logistic regression, Monocle’s Tobit model,^6^ DESeq2 1.16.11,^15^ MAST 1.2.1,^3^ and SCDE 1.99.4^2^ were used to benchmark the simulations. Monocle’s Tobit model method, DESEq2, and MAST were invoked using Seurat’s wrapper functionality through the function Seurat::FindMarkers.^18^ The method ‘glm’ which is loaded by default via the R native stats library was used to perform logistic regression, by using the parameter family = ‘binomial’.

The .fastq files output from the RSEM simulations were quantified using kallisto v0.43. tximport^19^ was used to aggregate transcript-level counts and abundances into gene-level counts and abundances prior to inputting into the various methods. For logistic regression, SCDE, and DESeq2, the gene counts were used as input. For Monocle and MAST, the TPM abundances were used as input. In order to afford each method its optimal input, normalization and filtering methods native to each method were used. We did not perform any additional normalization or filtering.

For each gene, logistic regression was performed using cluster labels as response variable and transcript counts as predictors. Any transcript that was not expressed in >90% cells was filtered from the logistic regression. A likelihood ratio test using a null model of a logistic regression on the cluster labels with no predictors was used to determine p-values.

To perform logistic regression using TCCs, we regressed on the TCCs that mapped unambiguously to each gene. For genes with more 70 unambiguously corresponding TCCs (dimensionality of TCCs >30% of total cells), the Kolmogorov-Smirnoff test was performed on each equivalence class independently to test whether the TCCs of the two clusters were derived from the same underlying distribution. The TCC-level p-values were then aggregated with Lancaster’s method, weighted by the mean count per TCC, to obtain gene-level p-values.^11^

We benchmarked the effects of three types of normalization on the logistic regression method: 1) DESeq2’s method of normalization using size factors, 2) TPM normalization and 3) no normalization, to ensure that the better performance by logistic regression is not due to differences in normalization. To apply DESeq2’s method, size factors were calculated based on the transcript counts using DESeq2::calculateSizeFactors, and the normalized counts per cell were obtained by dividing by the cell’s size factor. TPM normalization was obtained by using kallisto’s quantification outputs.

The code required to reproduce the analysis and figures in this data are available at https://github.com/pachterlab/NYMP_2018.

## Acknowledgments

We thank Nicolas Bray, Jase Gehring and Valentine Svensson for discussion and comments on the manuscript. Harold Pimentel assisted with the simulations. We thank Charlotte Soneson and Mark Robinson for questions that led to the clarification of GDE, DGE, DTU and DTE in Supplementary Figure 1.

